# Barcode identification for single cell genomics

**DOI:** 10.1101/136242

**Authors:** Akshay Tambe, Lior Pachter

## Abstract

Single-cell sequencing experiments use short DNA barcode ‘tags’ to identify reads that originate from the same cell. In order to recover single-cell information from such experiments, reads must be grouped based on their barcode tag, a crucial processing step that precedes other computations. However, this step can be difficult due to high rates of mismatch and deletion errors that can afflict barcodes. Here we present an approach to identify and error-correct barcodes by traversing the de Bruijn graph of circularized barcode k-mers. This allows for assignment of reads to consensus fingerprints constructed from k-mers, and we show that for single-cell RNA-Seq this improves the recovery of accurate single-cell transcriptome estimates.

## Availability and implementation

Freely available source code is available at Github: https://github.com/pachterlab/Sircel This Github repository also contains iPython notebooks to reproduce all analysis presented in this paper.

## Introduction

Tagging of sequencing reads with short DNA barcodes is a common experimental practice that enables a pooled sequencing library to be separated into biologically meaningful partitions. This technique is in the cornerstone of many single-cell sequencing experiments, where reads originating from individual cells are tagged with cell-specific barcodes; as such, the first step in any single-cell sequencing experiment involves separating reads by barcode to recover single-cell profiles (Svensson et al., 2017; Trapnell, 2015); (Klein et al., 2015). For example, in the Drop-Seq protocol, which is a popular microfluidic-based single-cell experimental platform, DNA barcodes are synthesized on a solid bead support, using split-and-pool DNA synthesis (Macosko et al., 2015). Similar split-and-pool barcoding strategies are used in other single-cell sequencing assays such as Seq-Well (Gierahn et al., 2017) and Split-seq (Rosenberg et al., 2017). One consequence of this synthetic technique is that deletion errors are extremely prevalent; by some estimates 25% of all barcode sequences observed contain at least one deletion (Macosko et al., 2015). Ignoring such errors can therefore dramatically lower the number of usable reads in a dataset, while incorrectly grouping reads together can confound single cell analysis.

Current approach to “barcode calling”, the process of grouping reads together by barcode, use simple heuristics to first identify barcodes that are likely to be uncorrupted, and then “error correct” remaining barcodes to increase yields. However the complex nature of errors, that unlike sequencing based error also include deletions, can lead to large number of discarded reads (reads that could not be assigned to a barcode) (Macosko et al., 2015). Additionally, some current approaches requires the approximate number of cells in the experiment be known beforehand, and in some experimental contexts such information is not easily obtained.

The problem of identifying true barcodes from among many sequences corrupted by mismatch and deletion errors seemingly requires a multiple sequence alignment, from which errors can be detected and corrected (Zorita, Cuscó, & Filion, 2015). However unlike standard biological sequence alignment settings, the single-cell barcode identification problem requires analysis of millions, if not billions of different sequences. On the other hand, the problem is constrained in that the sequences are short (barcodes are typically 10-16 bp long) and the length of each barcode is the same and known.

To circumvent the need for complete (and intractable) multiple sequence alignment, we rely on a k-mer based approach that is both fast and robust to error. Our method makes use of the idea of circularizing the sequences that are to be error corrected, and rather than pursuing a multiple sequence alignment approach, we instead borrow ideas from genome assembly. However unlike assembly methods developed for reconstructing circular genomes (Hunt, 2015) our use of circularization is merely a method for adding robustness to the k-mer fingerprinting of barcodes.

Our methods are implemented in software called SIRCEL whose input is a list of reads and which outputs the number and sequences of cell-barcodes from error-containing datasets in an unbiased manner. Our implementation is robust to insertion, deletion, and mismatch errors, and requires a minimal number of user-inputted parameters. The output is compatible with downstream single-cell analysis tools such as kallisto. (N. L. Bray, Pimentel, Melsted, & Pachter, 2016; Ntranos, Kamath, Zhang, Pachter, & Tse, 2016)

## Methods and algorithms

K-mer counting is a fast and well-established technique that has previously been used to dramatically speed up the assignment of reads to transcripts for RNA-seq (N. L. Bray et al., 2016; Patro, Mount, & Kingsford, 2014) and metagenomics(Schaeffer, Pimentel, Bray, Mellsted, & Pachter, 2015) and as such might be applicable to barcode calling. We reasoned that by counting k-mers we could rapidly identify error-free subsequences within the context of a larger error-containing read. (Li, 2015; Skums et al., 2012) The intuition behind our approach lies in the fact that while many copies of the same barcode may contain a different profile of errors, pairs of such barcodes may share some overlapping subsequence that is error free. However as the barcode errors are expected to be random, it is unlikely that several reads will share the exact same error pattern. As such, frequently occurring k-mers would arise from error-free regions of barcodes, while no overlap would be expected from error-prone k-mers. Similar reasoning has been previously used to rapidly detect and reject error-containing reads from RNA-seq and DNA assembly (Liu, Schroder, & Schmidt, 2013; Skums et al., 2012).

One difficulty associated with error-correcting barcodes using this technique lies in the fact that barcodes are typically very short: for example Drop-Seq barcodes are 12 base pairs long. Conversely in order for a k-mer counting approach to be feasible we must pick a moderately large value for *k*, typically *k = 9*. As a result there are many positions on a barcode where a single error would ensure that none of its k-mers are shared with an error-free barcode. To circumvent this problem we circularize the barcode sequences before counting k-mers; this ensures that barcodes containing a single mismatch error still share k-mers with the error-free sequence, independent of where the error occurred within the barcode (**Fig 1A)**. Furthermore this approach allows for addressing the possibility of insertion or deletion errors. When circularizing the barcode sequence, we can first either extend or truncate the sequence by one nucleotide. Doing this provides the same robustness to positional errors, but additionally allows for robustness to insertion or deletion errors. As every read contains unknown mutation type(s), we perform all three circularization operations before counting k-mers. Thus, we obtain a set of error-free subsequences that derive from the ‘true’ barcodes. This procedure guarantees that all reads with either zero or one error contribute some error-free k-mers, while reads with two or more errors sometimes contribute error-free k-mers, depending on the spacing between the errors.

**Figure 1.**
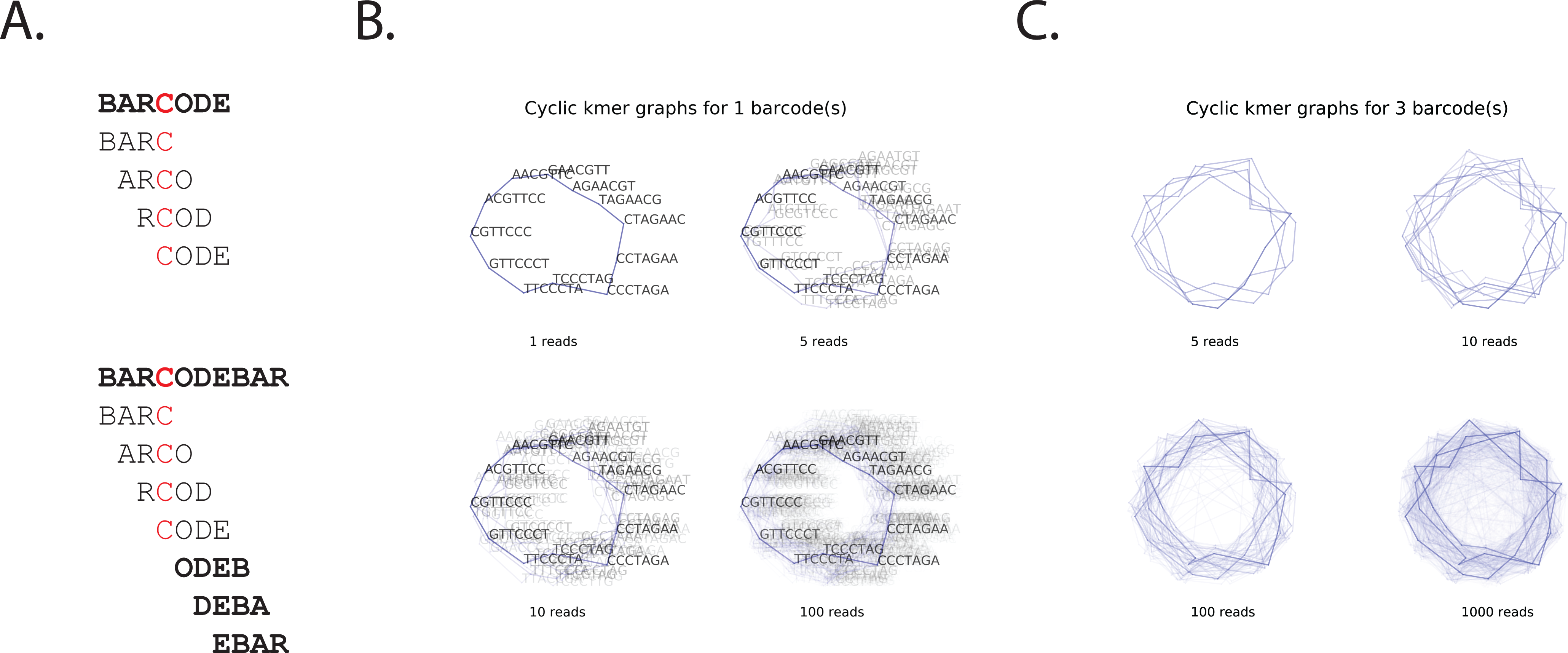
A strategy to use k-mer counting to identify sequence barcodes. A. Circularizing barcodes ensures robustness against single mismatches. An example sequence ‘BARCODE’ contains an error (highlighted in red). When the barcode sequence is short relative to k, all k-mers from this sequence will contain the mutated base. Circularizing the sequence (bottom) ensures that there will be some error-free k-mers from a sequence independent of the position of the error. B. An example circular k-mer graph containing one barcode. Error-containing reads were simulated from a ground-truth barcode. Reads were circularized and k-mers were counted. The resultant k-mer graph is plotted here. Nodes in this graph are represented as gray dots, and edges as blue lines. Edges weights are represented by shading (dark = high edge weight). Despite a fairly high rate of error (Poisson 3 errors per 12 nucleotide barcode), the true barcode path is visually discernable with a modest number of reads. C. An example circular k-mer graph containing three barcodes. Same as above.

We use these k-mer counts to identify and error-correct complete barcodes. To do this we build and traverse a directed, weighted de Bruijn graph (Compeau, Pevzner, & Tesler, 2011). In this graph, nodes represent subsequences of length *k - 1*, and an edge represents two nodes that directly adjacent to each other in at least one k-mer. The weight of these edges relates to how many times each edge (k-mer) was observed in the entire dataset. Additionally as the barcode portions of these reads are stranded, these edges are directed by the order of their appearance in the read (5’ to 3’). In this graph, which originates from circularized barcode sequences, a cyclic path of length *l* represents a possible barcode sequence of the same length. We define the capacity of a path to be the weight of the lowest edge within that path. Thus, high-weight paths represent possible barcodes that contain frequently observed k-mers, while low-weight paths likely represent cycles that formed spuriously. This is depicted in Fig 1B and 1C. We emphasize here that we do not need any single read to contain all k-mers in a high-weight path / error-corrected barcode; it is the overlap of many k-mers that likely originate from a number of reads that gives rise to such a path.

To rapidly identify cyclic paths from this graph we use a greedy depth-first recursive search (Algorithm 1). Briefly, this algorithm works by first (randomly) picking a node from the graph to initialize the search. Each of the outgoing edges that connect to this node is checked, in descending order of the edge weights. This step is repeated for each of the children nodes, for a fixed number of steps given by the length of the barcode (a user-suppled parameter). If at the end of these steps the procedure returns to the same node where it began, a cycle has been found. This is described in some more detail in Algorithm 1.

This approach identifies several cyclic paths from the barcode de Bruijn graph, and the depth of this search is determined by user-supplied parameters. As only a subset of these paths represents a true error-corrected barcode sequence, we filter the paths based on their path weight. We hypothesized that paths representing a true error-corrected barcodes would have a higher capacity than paths that contained errors, or paths formed by spurious k-mer overlap between [barcode-wise] unrelated sequences. To verify this hypothesis we plotted the cumulative distribution of path capacities, and observed a clear inflection point, corresponding to a subset of paths that had a significantly higher capacity than the rest of the population. We computationally identified this inflection point as a local maximum in the first derivative of the cumulative distribution function. This was facilitated by first smoothing the CDF.

Paths are then thresholded at this inflection point, and paths with capacities higher than the threshold value are deemed error-corrected barcodes, while the rest of the paths are rejected. We then assign each read in our dataset to one of the error-corrected barcodes based on k-mer compatibility. in other words, a read is assigned to the consensus barcode with which it shares the most k-mers. At this point we can vary the value of $k$. Preparing the barcode de Bruijn graph with a larger *k* than that used to assign reads enables us to call error-free barcodes with the higher stringency, while assigning a large number of reads (Algorithm 2).

Finally, to improve performance we make a small modification to the protocol outlined above. Rather than building a de Bruijn graph of the entire barcode dataset, we instead build a new subraph for each new node we initialize the search with. This subgraph contains all nodes that are indirectly connected (within a fixed number of steps) to the node at which we initialize the search. This substantially simplifies the search procedure while leaving performance unaffected. To rapidly build these subgraphs we prepare a k-mer index of the input dataset, which maps a k-mer to a list of reads that contains that k-mer. When performing a search from a random start node, we query the k-mer index for the start node and prepare a de Bruijn graph from only the subset of reads returned by the query.

As this index can be quite large (for Drop-seq, which uses 12mer barcodes, each read produces 36 circularized and truncated / extended k-mers to be indexed), which results in an extremely large index. We further simplify this protocol by preparing the index from a subset of the reads. This approximation also does not affect performance, as long as the subset is ‘representative’ of the entire dataset. The exact parameters for this depend on the sequencing depth, number of barcodes, error rate and likely other parameters; however in our tests simply indexing ∼1m reads is sufficient (Table 1).

**Table 1.** Run time for downsampled Macosko et al., datasets

| Number of reads in dataset | Number of reads indexed | Number of cells detected | Time |
| --- | --- | --- | --- |
| 1,000,000 | 100,000 | 562 | 6m 39s |
| 1,000,000 | 1,000,000 | 582 | 14m 57s |
| 10,000,000 | 100,000 | 575 | 42m 88s |
| 10,000,000 | 1,000,000 | 584 | 48m 55s |
| 100,000,000 | 100,000 | 574 | 154m 11s |
| 100,000,000 | 1,000,000 | 585 | 207m 57s |

## Results

We validated our approach by attempting to identify and error-correct barcode sequences in both real and simulated datasets. We re-analyzed a previously published species-mixing Drop-Seq experiment published by Macosko et al. This experiment involved single-cell sequencing of a mixture of human and mouse cells, and as such it served as a useful control for barcode calling: if the calling performs well, cells should only contain human, or mouse reads but not both. We used our algorithm on this data and as expected found a clear inflection point in the cumulative distribution of barcode paths. We could readily identify this inflection by its smoothed first derivative, and thresholding the paths at this inflection point yielded 582 barcodes, each of which had accounted for approximately the same number of reads (Fig. 2A and 2B). These values were consistent with previously reported values from the same dataset. We then quantified single-cell expression profiles using combined human / mouse transcriptome, once again using an algorithm derived from k-mer counting [kallisto]. As seen in Fig. 2C, ‘cells’ that consist of reads clustered by similar barcode k-mers also exhibit distinct expression profiles. In nearly every case cells appear to consist of reads deriving entirely from one species (Fig. 2D). This result indicates that our k-mer counting approach can be used to group reads into single-cell datasets.

**Figure 2.**
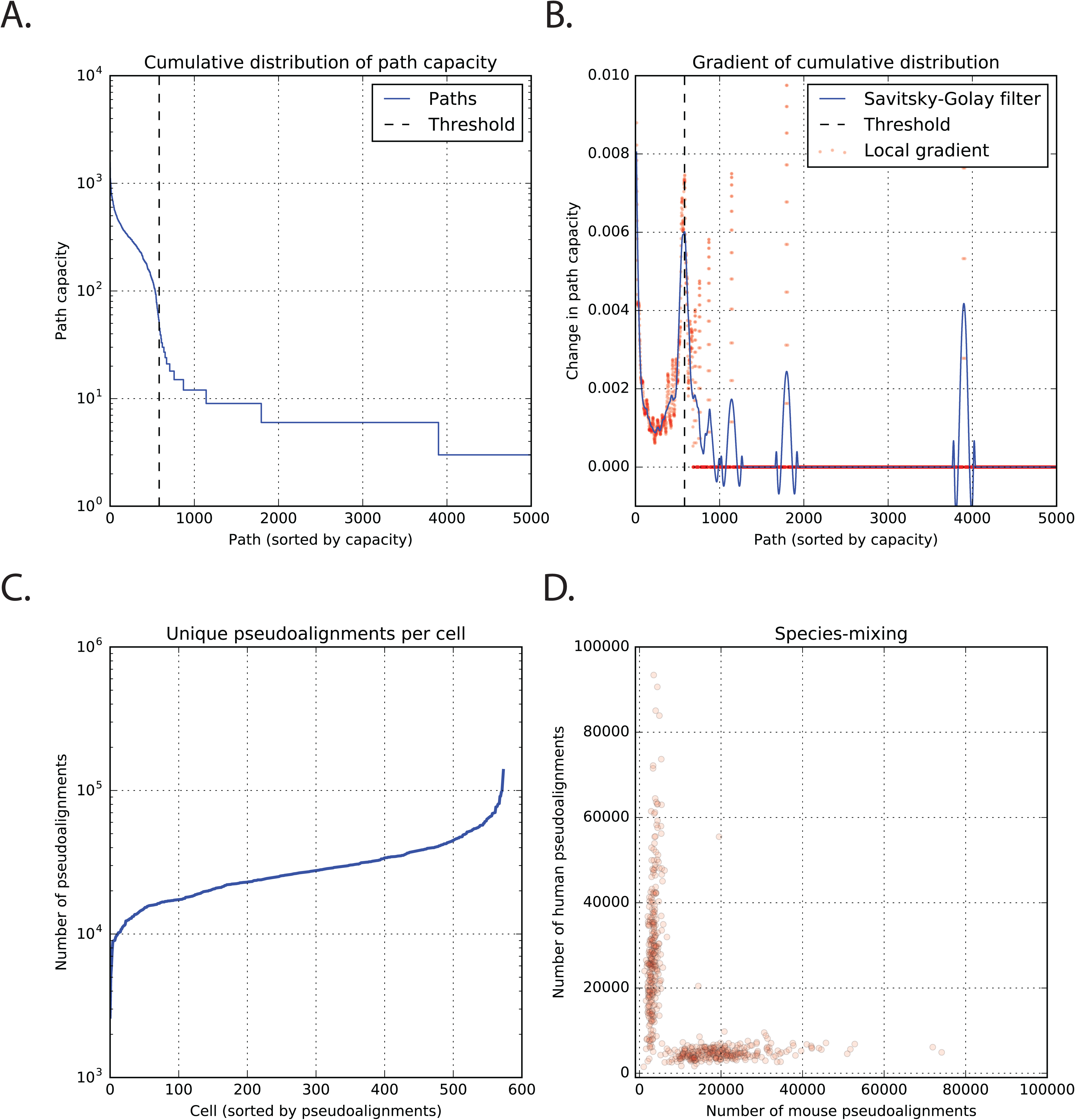
Identifying barcodes and splitting reads from Macosko et al., species mixing experiment. A. Circular paths were identified in the circular barcode k-mer graph from a published Drop-seq dataset. The cumulative distribution of circular path weights (shown here) clearly shows an inflection point. Paths with weight higher than this inflection point are deemed to be true barcodes. B. This inflection point can be identified as a local maximum in the first derivative of the cumulative distribution. A Savitskiy Golay filter facilitates in this identification by smoothing the data. C. Reads were grouped into cells by assigning them to to thresholded paths based on k-mer compatibility alone. This assignment results in a flat distribution in the number of pseudoalignments per cell. D. Reads that were split based on barcode k-mer compatibility alone also segregate by their number of pseudoalignments to different transcriptiomes. This indicates that assigning reads based on k-mer compatibility produces distinct and biologically relevant groupings.

To benchmark our algorithms’ performance and establish it’s performance limits, we performed a large number of simulations, under a wide variety of scenarios. We produced a fixed number of ‘true’ barcodes, and produced reads by adding a Poisson number of errors to each read. Error positions were selected uniformly at random, and separate datasets were produced for insertion, deletion, mismatch, and all errors. We also varied the barcode abundance distributions between normal, uniform and exponential. For each condition we produced 3 separate datasets and evaluated our algorithms’ performance on each. As shown in **Fig S1**, our algorithm is able to identify the error-free barcode sequences independent of the number of Poisson errors per read. However we do see a dependence on the specific error type: mismatch errors are better tolerated than insertion or deletion errors. Additionally we find that the barcode abundance distribution strongly affects our ability to detect and error correct barcodes, especially as the error rate increasese.

We also used our simulations to evaluate how well we could assign reads to error-corrected barcodes. From each true positive barcode detected from our simulations, we computed the fraction of reads that were correctly assigned to its consensus barcode. The median value of this parameter is shown in **Fig S2**, as a function of Poisson error rate, and the distribution of this value over all cell barcodes is shown in **Fig S3**. Here we observe a similar trend: whether or not a read is correctly assigned depends on the error type, as well as barcode abundance distribution. Once again mismatch errors are better tolerated than either insertions or deletions, while exponentially distributed barcode abundances are tolerated worse than normally or uniformly distributed barcodes.

To evaluate the extent to which indexing a subset of reads affects the ability to call barcode clusters, we systematically subsampled the raw data from Macosko et al., and ran our software on these datasets. For each subsampled dataset we indexed either 100,000 or 1,000,000 reads. As seen in Table 1, the number of cells detected is essentially independent of either the number of reads in the dataset, and the number of reads indexed, showing that indexing ∼1m reads is sufficient to error-correct read barcodes. Furthermore we point out that with this indexing strategy the runtime to index a dataset and detect consensus barcodes is constant and independent of the number of reads in the dataset. The runtime to assign reads to consensus barcodes increases linearly with the number of reads (Table 1).

## Discussion

We have shown how a de Bruijn graph formulation of the barcode calling problem based on circularization of input sequences is a useful approach to identify and error-correct barcode sequences. Our approach simplifies the problem of sequence error correction by rephrasing it as a k-mer counting question, and as such is simple and relatively fast. Furthermore it does not rely heavily on user-supplied parameters or any prior knowledge about the exact nature of the sequencing errors; as such we expect it to be applicable to a number of different single-cell barcoding techniques that differ in the exact nature of the barcode generation chemistry. We also show that our approach produces usable data from real-world datasets, and that our integrated pipeline using kallisto and transcript compatibility counts is an effective approach for rapid and accurate analysis of Drop-Seq single-cell RNA-Seq data.

We benchmarked our algorithm using an extensive set of simulations that systematically varied the error rate per read, the error type, and the abundance of each barcode [single-cell] within the dataset. From these simulations we observe that the barcode abundance distribution makes a significant difference to performance, with normally- and uniformly- distributed barcode abundances being far better tolerated than exponentially distributed barcodes. This behavior is expected; with exponentially distributed barcode abundances, the inflection point in the CDF of cyclic path weights is obscured, making it difficult to distinguish between a true barcode path with low abundance, and an error-containing path with relatively high weight. Notably, exponentially distributed barcode abundances are not expected (and indeed not observed) in real data: the total RNA content from any given single cell in an experiment are typically approximately uniformly distributed.

These simulations also demonstrated that although our method is tolerant of errors when identifying and error-correcting barcode sequences, errors lower its ability to assign individual reads to error-corrected consensus sequences. This is not surprising, because reads with a large number of errors are unlikely to contain any error-free k-mers that are required to assign a read. This effect can be mitigated by using a smaller value of *k* when assigning reads. Simulations also revealed that our algorithm is more tolerant to mismatch errors over insertion or deletion errors. We postulate that this is because in the barcode de Bruijn graph, reads that contain only mismatches form a cyclic path of the correct length, whereas reads containing insertion or deletion errors form paths with incorrect length, complicating the cyclic-path search protocol. This effect is most pronounced at the error rates that are higher than typical Drop-seq datasets.

## Conclusion

Single-cell genomics is a dynamic field that encompasses a large and growing number of techniques that measure a variety of biological properties. However one commonality in these workflows is that experiments mark reads originating from distinct cells with single cell barcodes. Correctly identifying and grouping reads by their barcodes in the presence of experimental and sequencing errors is an essential first step in any single-cell analysis pipeline. The software presented here addresses this universal problem, and as such it should be useful for a variety of single cell sequencing based genomics experiments. Although we focus here on the specific Drop-seq protocol, there are a number of related single-cell experiments that rely on split-pool combinatorial synthesis of barcodes (Rosenberg et al., 2017), as well as other massively parallel single-cell sequencing experiments that measure other genomic and transcriptomic properties (Gierahn et al., 2017; Rotem et al., 2015). As error correcting and clustering barcodes is central to these assays as well, we believe that these methods will also benefit from our software.

## Acknowledgements

We thank Jase Gehring and Vasilis Ntranos for helpful comments and feedback during the development of the method.

## Example

The example data set (supplementary data) shows the workflow to identify and split barcoded reads from a published Drop-seq dataset [SRR1873277]. This dataset derives from a species-mixing experiment, where human and mouse cells were mixed prior to single-cell RNA-seq. As such reads grouped by their barcodes should also segregate by which species they [pseudo]align with. We can therefore evaluate the performance of Sircel by how frequently reads from the two species appear to derive from the same cell.

## Methods

Raw sequencing data for Macosko et al. species mixing was obtained from the sequence read archive (SRR1873277) and converted to fastq format using SRA-toolkit. Subsamples of these datasets were obtained using standard command line tools:

~~~
zcat INFILE.fastq.gz | head –n NUM_READS*4 | gzip > OUTFILE.fastq.gz
~~~

We used this data without any further processing, or read filtering. Sircel was then used to identify barcodes and assign reads with the following parameters: k-mer length of 7, search breadth of 1000 subgraphs, search depth of 5 paths per subgraph. All results presented here were processed with 32 threads.Output from Sircel was then fed, into a single-cell analysis pipeline based on kallisto (N. L. Bray et al., 2016)and transcript compatibility counts (Ntranos et al., 2016). Our integrated pipeline, as well as ipython notebooks to visualize the data is available on Github.

Simulations were performed by first randomly generating a 500 ground truth barcode sequences of length 12. Each barcode was assigned a relative abundance drawn from one of three pre-defined distributions (normal, uniform and exponential). Reads were generating reads by selecting a barcode according to the barcode abundance, and adding a Poisson number of errors given by user-defined rate. Error type (insertion, deletion, mismatch or any) was also varied systematically during this step. Each simulation consisted of 100,000 reads generated in this manner. For each condition (combination of barcode abundance distribution, Poisson error rate and error type), we produced three separate simulations for a total of 180 datasets.

These simulated datasets were then fed into Sircel to identify error-free barcodes. For each simulation we compared the output of Sircel to the ground-truth barcodes, identifying true positives as barcode sequences that were present in both the Sircel output and the ground-truth, false positives as barcode sequences that were found in the Sircel output but not the ground truth, and false negatives as barcode sequences that were not found in the Sircel output but not the ground truth. For each true positive barcode identified by Sircel, we additionally evaluated whether the reads assigned to that barcode were correctly assigned. Reads that derived from the ground truth barcode were deemed correctly assigned, and all other reads were labeled as incorrectly assigned.

Ipython notebooks to reproduce this analysis are available on Github.

### Algorithm 1. Recursively identify cycles of fixed in graph

1. Initialize recursion:

a. Pick a starting edge

i. Edge links node and neighbor

1. Set the current path to the starting edge
2. Record the identity of the starting node
2. Recursion:

a. Get all outgoing edges that emanate from neighbor
b. Sort outgoing edges by edge weight (descending)
c. For each outgoing edge

i. Extend the current path by this edge
ii. Continue recursion
3. Terminate recursion:

a. If the current path length is longer than the barcode length

i. If the path start node and end node are the same, the path is a cycle

1. Return True
ii. Else return False

### Algorithm 2. Identify barcodes and assign reads

1. Index k-mers

a. Build a look-up table associating each k-mer with a set of reads that contains that k-mer
2. Identify barcodes

a. Pick a starting edge (k-mer) in order by edge weight
b. Build a de Bruijn sub-graph. This is a directed, weighted, de Bruijn graph containing only the subset of reads that contain the starting k-mer

i. Building only a subgraph greatly speeds up graph traversal
c. Identify high-weight cycles

i. **See** Algorithm 1.
ii. Remove this cycle from the subgraph by decrementing the weights of all edges in this subgraph by the capacity of this path
d. Repeat c to find other paths that originate from this node
e. Repeat a–e to find paths that start at a different node
3. Merge similar paths by Hamming distance
4. Threshold paths to identify true barcodes

a. Identify an inflection point in the cumulative distribution of path lengths
5. Assign reads

a. Assign each read to the path with which it shares the most k-mers

---

**Figure S1. Identifying consensus barcodes is robust to a high number of errors per barcoded, but sensitive to the distribution of barcode abundances**

We performed several simulations with error-prone reads deriving from 500 randomly generated barcode sequences. The number of errors per read, the type of errors, and the distribution of barcode abundances were all systematically varied. Performance was evaluated by quantifying the number of true positive consensus barcodes (blue), false positive barcodes (red) and false negative barcodes (yellow). Data shown here represents 5 separate simulations.

A. Normally distributed barcode abundances.

B. Uniformly distributed barcode abundances.

C. Exponentially distributed barcode abundances

**Figure S2. Assigning reads to consensus barcodes by k-mer compatibility depends on errors rate and barcode abundance distribution**

We performed several simulations with error-prone reads deriving from 500 randomly generated barcode sequences. The number of errors per read, the type of errors, and the distribution of barcode abundances were all systematically varied. The fraction of reads that were correctly assigned in each cell was quantified, and the median value for this parameter in each detected cell is shown here, as a function of Poisson error rate. Data shown here represents 5 separate simulations.

A. Normally distributed barcode abundances.

B. Uniformly distributed barcode abundances.

C. Exponentially distributed barcode abundances

**Figure S3. Assigning reads to consensus barcodes by k-mer compatibility depends on errors rate and barcode abundance distribution**

We performed several simulations with error-prone reads deriving from 500 randomly generated barcode sequences. The number of errors per read, the type of errors, and the distribution of barcode abundances were all systematically varied. The fraction of reads that were correctly assigned in each cell was quantified, and the distribution of this value over all cells is shown here. Data shown here represents 5 separate simulations.

A.Normally distributed barcode abundances with 1 error per read

B. Uniformly distributed barcode abundances with 1 error per read

C. Exponentially distributed barcode abundances with 1 error per read

D. Normally distributed barcode abundances with 2 errors per read

E. Uniformly distributed barcode abundances with 2 errors per read

F. Exponentially distributed barcode abundances with 2 errors per read

**Figure S4. Circularized de Bruijn graph from real data**.

A de Bruijn subgraph was prepared from circularized reads that could be assigned assigned to 10 randomly selected barcodes from the Macosko et al dataset is depicted here. Line transparency is proportional to the weight of each edge.

